# Transcriptomic atlas of mushroom development highlights an independent origin of complex multicellularity

**DOI:** 10.1101/349894

**Authors:** Krisztina Krizsán, Éva Almási, Zsolt Merényi, Neha Sahu, Máté Virágh, Tamás Kószó, Stephen Mondo, Brigitta Kiss, Balázs Bálint, Ursula Kües, Kerrie Barry, Judit Cseklye, Botond Hegedűs, Bernard Henrissat, Jenifer Johnson, Anna Lipzen, Robin A. Ohm, István Nagy, Jasmyn Pangilinan, Juying Yan, Yi Xiong, Igor V. Grigoriev, David S. Hibbett, László G. Nagy

**Affiliations:** Synthetic and Systems Biology Unit, Institute of Biochemistry, BRC-HAS, Szeged, 6726, Hungary.; US Department of Energy (DOE) Joint Genome Institute, Walnut Creek, CA, 94598, United States.; Seqomics Ltd. Mórahalom, Mórahalom 6782, Hungary; Division of Molecular Wood Biotechnology and Technical Mycology, Büsgen-Institute, University of Göttingen, Göttingen, Germany.; Institute of Biophysics, BRC-HAS, Szeged, 6726, Hungary.; Architecture et Fonction des Macromolécules Biologiques (AFMB), UMR 7257 CNRS Université Aix-Marseille, 13288, Marseille, France.; Department of Biology, Microbiology, Utrecht University, 3584 Utrecht, The Netherlands; Biology Department, Clark University, 01610 Worcester MA, USA

## Abstract

We constructed a reference atlas of mushroom formation based on developmental transcriptome data of six species and comparisons of >200 whole genomes, to elucidate the core genetic program of complex multicellularity and fruiting body development in mushroom-forming fungi (Agaricomycetes). Nearly 300 conserved gene families and >70 functional groups contained developmentally regulated genes from five to six species, covering functions related to fungal cell wall (FCW) remodeling, targeted protein degradation, signal transduction, adhesion and small secreted proteins (including effector-like orphan genes). Several of these families, including F-box proteins, protein kinases and cadherin-like proteins, showed massive expansions in Agaricomycetes, with many convergently expanded in multicellular plants and/or animals too, reflecting broad genetic convergence among independently evolved complex multicellular lineages. This study provides a novel entry point to studying mushroom development and complex multicellularity in one of the largest clades of complex eukaryotic organisms.

---

Mushroom-forming fungi (Agaricomycetes, Basidiomycota) represent an independent lineage of complex multicellular organisms with a unique evolutionary history compared to complex animals, plants and stramenopiles. They comprise >21,000 species and originated ~350 million years ago^1^, approximately coinciding with the origin of tetrapods. Mushrooms have immense importance in agriculture, ecology and medicine; they represent an important and sustainable food source, with favorable medicinal properties (e.g. antitumor, immunomodulatory)^2^.

Complex multicellular development in fungi has been subject to surprisingly few studies^3–6^, resulting in a paucity of information on the genetic underpinnings of the independent origins of complex multicellularity in fungi^7^. As complex multicellular structures, fruiting bodies deploy mechanisms for hypha-to-hypha adhesion, communication, cell differentiation, defense and execute a developmental program that results in a genetically determined shape and size^6,7^. Fruiting bodies shelter and protect reproductive cells and facilitate spore dispersal. Uniquely, complex multicellularity in fungi comprises short-lived reproductive organs whereas in animals and plants it comprises the reproducing individual. Nevertheless, fruiting bodies evolved complexity levels comparable to that of simple animals, with up to 30 morphologically distinguishable cell types described so far. Fruiting body development is triggered by changing environmental variables (e.g. nutrient availability), and involves a transition from simple multicellular hyphae to a complex multicellular fruiting body initial. The initial follows genetically encoded programs to develop species-specific morphologies^5,6^, which in the Agaricomycetes ranges from simple crust-like forms (e.g. *Phanerochaete*) to the most complex toadstools (e.g. *Agaricus bisporus*). Previous studies identified several developmental genes, including hydrophobins^8^, defense-related proteins^9^, fungal cell wall (FCW) modifying enzymes^10–13^, transcriptional regulators^4,5,14^ (e.g. mating genes) and light receptors^15^ (e.g white collar complex). It is not known, however, what genes comprise the ‘core toolkit’ of multicellularity and development in the Agaricomycetes.

Here we investigate the general evolutionary and functional properties of fruiting body development and assess whether mushroom-forming fungi evolved complex multicellularity via unique or convergent solutions, compared to other complex lineages (e.g. plants, animals or brown algae). We combine comparative analyses of developmental transcriptomes of six species with comparisons of 201 whole genomes and focus on conserved developmental functions and complex multicellularity.

## Results

We obtained fruiting bodies in the laboratory or from the field and profiled gene expression in 3-9 developmental stages and tissue types for *Coprinopsis cinerea* AmutBmut, *Schizophyllum commune* H4-8, *Phanerochaete chrysosporium* RP78, *Rickenella mellea* SZMC22713 and *Lentinus tigrinus* RLG9953-sp. We used data for *Armillaria ostoyae* C18/9 from our previous work^16^. We report the *de novo* draft genome of *Rickenella mellea* (Hymenochaetales), the phylogenetically most distant species from *Coprinopsis* in our dataset, spanning >200 million years of evolution^1^. To construct a reference atlas of mushroom development, we performed poly(A)+ RNA-Seq on Illumina platforms, in triplicates (totalling to >120 libraries, Supplementary Table 1). We obtained an average of 60.8 million reads per sample, of which on average 83.3% mapped to the genomes (Supplementary Fig 1). For each species, the first and last developmental stage sampled were vegetative mycelium and mature fruiting body at the time of spore release, respectively. This spans all developmental events of fruiting bodies except senescence. Because it is difficult to align intermediate developmental stages across species, we identified developmentally regulated genes using an approach that removes developmental stage identity from the analyses (see Methods). Using this strategy, we could recover >80% of previously reported developmental genes of *Coprinopsis* (Supplementary Table 2). To more broadly infer functionalities enriched in mushroom forming fungi, we analyzed Interpro domain counts across 201 fungal genomes (including 104 Agaricomycetes), which revealed 631 significantly overrepresented domains in mushroom-forming fungi (*P* < 0.01, Fisher exact test, abbreviated FET, Supplementary Table 3-4).

### Dynamic reprogramming of the fungal transcriptome

We detected 12,003 - 17,822 expressed genes, of which 938-7,605 were developmentally regulated in the six species (Fig. 1/a, Supplementary Table 5). We found 192-7,584 genes that showed significant expression dynamics during fruiting body development (‘FB’ genes). Of developmentally regulated genes 188-1,856 genes were upregulated at fruiting body initiation (‘FB-init’ genes), which represents a transition from simple to complex multicellular organization. Only *Phanerochaete chrysosporium* had more FB-init genes than FB genes, which is consistent with its fruiting bodies being among the least complex types in the Agaricomycetes. The number of genes significantly differentially expressed (DEG) at fruiting body initiation further suggests that the transition to complex multicellularity is associated with a major reprogramming of gene expression (Supplementary Fig. 2). The largest numbers of DEGs were observed in cap and gill tissues in all four species with complex fruiting bodies. On the other hand, the expression profiles of stipes changed little relative to primordium stages of *Armillaria, Lentinus* and *Rickenella*, which is explained by the differentiation of the cap initial at the top of a primordial stipe, as opposed to *Coprinopsis*, in which stipe and cap differentiation happens simultaneously inside the fruiting body initial^17^. Many Gene Ontology (GO) terms were partitioned between vegetative mycelium and fruiting body samples (P<0.05, FET). Terms related to fungal cell wall, oxidoreductase activity and carbohydrate metabolism were enriched in developmentally regulated genes of all six species (Fig. 1/b, Supplementary Table 6), suggesting that cell wall remodeling is a common upregulated function in fruiting bodies. Other commonly enriched terms cover functions such as DNA replication, transmembrane sugar transport, ribosome, membrane and lipid biosynthesis, while many other were specific to single species (Supplementary Fig. 3).

**Fig. 1.**
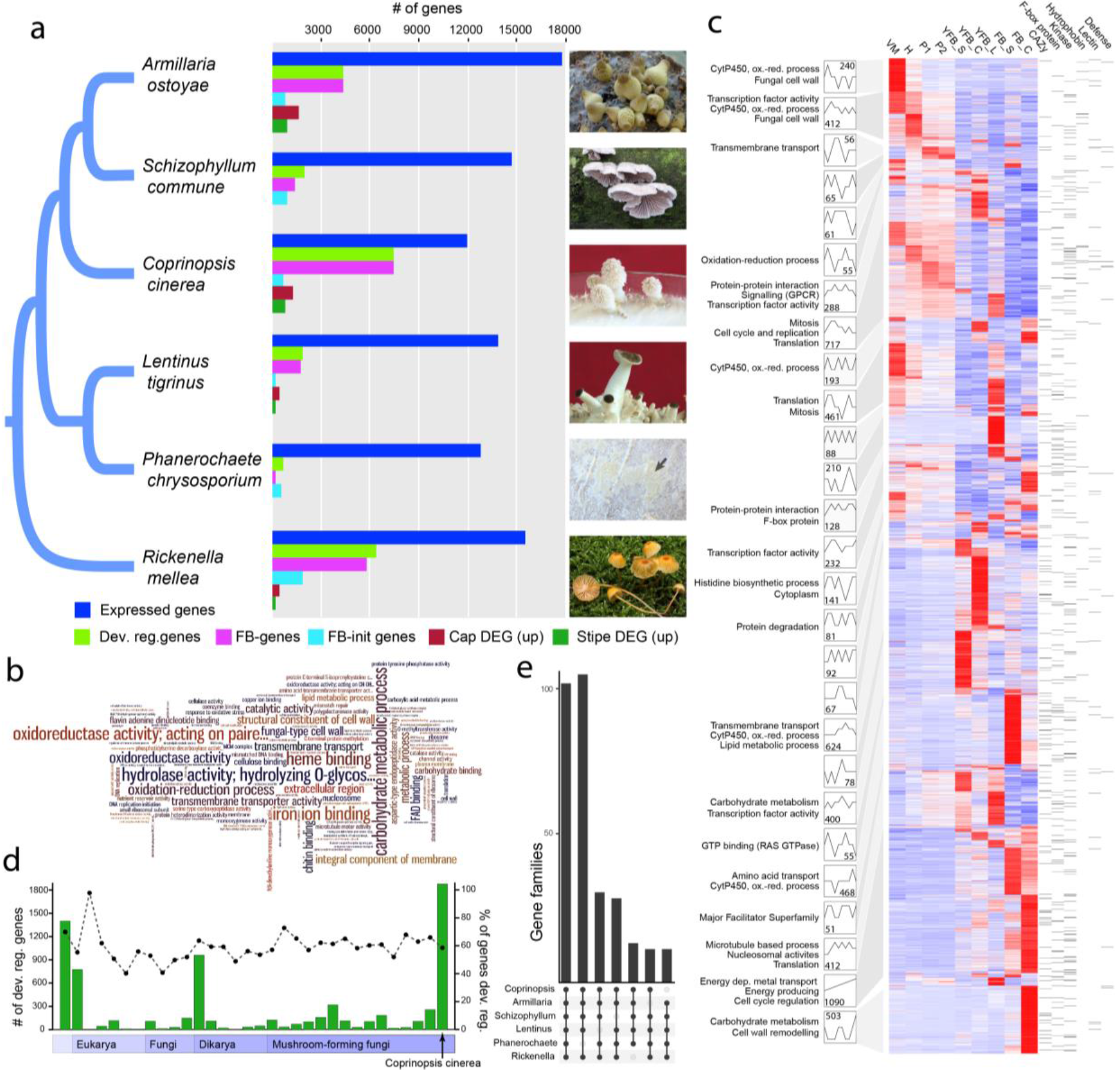
Overview of the developmental transcriptomes, **a**, the distribution of developmentally regulated and cap and stipe differentially expressed (DEG) genes in six mushroom-forming species. Fruiting bodies are shown on the right (*Phanerochaete*: K. Krizsán, *Rickenella:* Bálint Dima, others: L.G. Nagy). Two groups of developmentally regulated genes were defined: those that show >4 fold change and a FPKM>4 between any two stages of fruiting body development (referred to as ‘FB genes’) and that show >4 fold increase in expression from vegetative mycelium to the first primordium stage (referred to as ‘FB-init genes’). These definitions exclude genes that show highest expression in vegetative mycelium and little or no dynamics later on. **b**, wordcloud of Gene Ontology terms enriched in the developmentally regulated genes in the six species. Word size corresponds to the number of species in which a term was significantly enriched (*P* < 0.05, Fisher exact test). **c**, Analysis of co-expression modules in *Coprinopsis cinerea.* Heatmap of 7475 developmentally expressed genes arranged based on module assignment, with simplified expression profiles and the GO enrichment terms given for each module (see also Supplementary Table 7). We graphically depict only 27 modules with >50 genes (refer to Supplementary Fig. 4 for the complete list and data for other species). The distribution of key developmental genes is given on the right side of the heatmap. **d**, Phylostratigraphic profile of developmentally regulated genes of *Coprinopsis cinerea.* Genes are divided into age categories from genes shared by all cellular life (left) to species-specific genes (right). The percentage of developmentally regulated genes compared to all genes in a category is shown by a dashed line. Data for other species is presented in Supplementary Fig. 5. **e**, UpSetR representation of gene families developmentally regulated in at least 5 species. See also Supplementary Fig. 6.

To obtain a higher resolution of developmental events, we arranged developmentally regulated genes into co-expression modules using the Short Time Series Expression Miner (STEM)^18^. Developmentally regulated genes grouped into 28-40 modules, except *Phanerochaete* which had eleven. The largest modules in all species contained genes expressed at fruiting body initiation or in early primordia and genes with tissue-specific expression peaks, in young fruiting body caps, gills, stipes, mature fruiting bodies and stipes or caps (Fig 1/c; for further data see Supplementary Fig. 4 and Supplementary Note 1). Many early-expressed modules show upregulation across multiple stages (hyphal knot, stage 1 and 2 primordia), suggestive of an early expression program overarching multiple primordium stages. Co-expression modules display distinct functional enrichment signatures, as shown on Fig 1/c and Supplementary Table 7. For example, DNA replication and mitosis were characteristic for early-expressed modules, consistent with an early wave of nuclear and cellular division events followed by cell expansion without significant change in cell numbers ^5^. Growth by cell expansion is a mechanism shared with plants and possibly reflects constraints imposed by independently evolved rigid cell walls in these groups.

### Most developmental gene families are older than fruiting body formation

We investigated the evolutionary age distribution of developmentally regulated genes using phylostratigraphy^19^, based on a dataset of 4,483 archaeal, bacterial and eukaryotic genomes^20^ supplemented with 416 fungi (of which 113 were Agaricomycetes). We assigned genes to phylogenetic ages, ‘phylostrata’, by identifying for each gene the most phylogenetically distant species in which a homolog could be detected. The phylostratigraphic profiles of all species show three peaks, corresponding to two major periods of fungal gene origin: the first containing genes shared by all living species, the second by the Dikarya (Ascomycota + Basidiomycota) and a third containing species-specific genes. Many developmentally regulated genes have homologs in simple multicellular and unicellular eukaryotes and prokaryotes: the origin of 49.3 - 63.3% predate the origin of mushroom-forming fungi (Fig. 1/d, Supplementary Fig. 5), indicating that co-option of conserved genes contributed significantly to the evolution of fruiting bodies. Nevertheless, Agaricomycetes-specific phylostrata showed a characteristic enrichment for F-box genes, transcription factors and protein kinases, indicating an increased rate of origin for these in mushroom-forming fungi (Supplementary Table 8).

### Splicing patterns associate with development

We reconstructed transcript isoforms across developmental stages and tissue types in the six species using region restricted probabilistic modeling, a strategy developed for gene-dense fungal genomes^21^. We found evidence of alternative splicing for 36-46% of the expressed genes (Supplementary Table 9), which is significantly higher than what was reported for fungi outside the Agaricomycetes^22,23^ (1-8%). This transcript diversity was generated by 6,414 - 13,780 splicing events in the six species. Of the four main types of events, intron retention (44.3-60.5%) was the most abundant in all species, followed by alternative 3’ splice site (22.9-30.1%), alternative 5’ SS (15.6-24.1%) and exon skipping (0.8-2.9%) (Fig. 2/a), consistent with observations made on other fungi^22–24^. No substantial difference in the proportion of spliced genes and of splicing events was observed across developmental stages, tissue types or species. Nevertheless, we found that several genes with nearly constant overall expression level had developmentally regulated transcript isoforms (Fig 2/b-c). The six species had 159-1,278 such genes, the highest number in *Rickenella* (1,278) and the lowest in *Phanerochaete* (159) (Fig 2/d, Supplementary Table 9). Based on their expression dynamics, these transcripts potentially also contribute to development, expanding the space of developmentally regulated genes through alternative splicing.

**Fig. 2.**
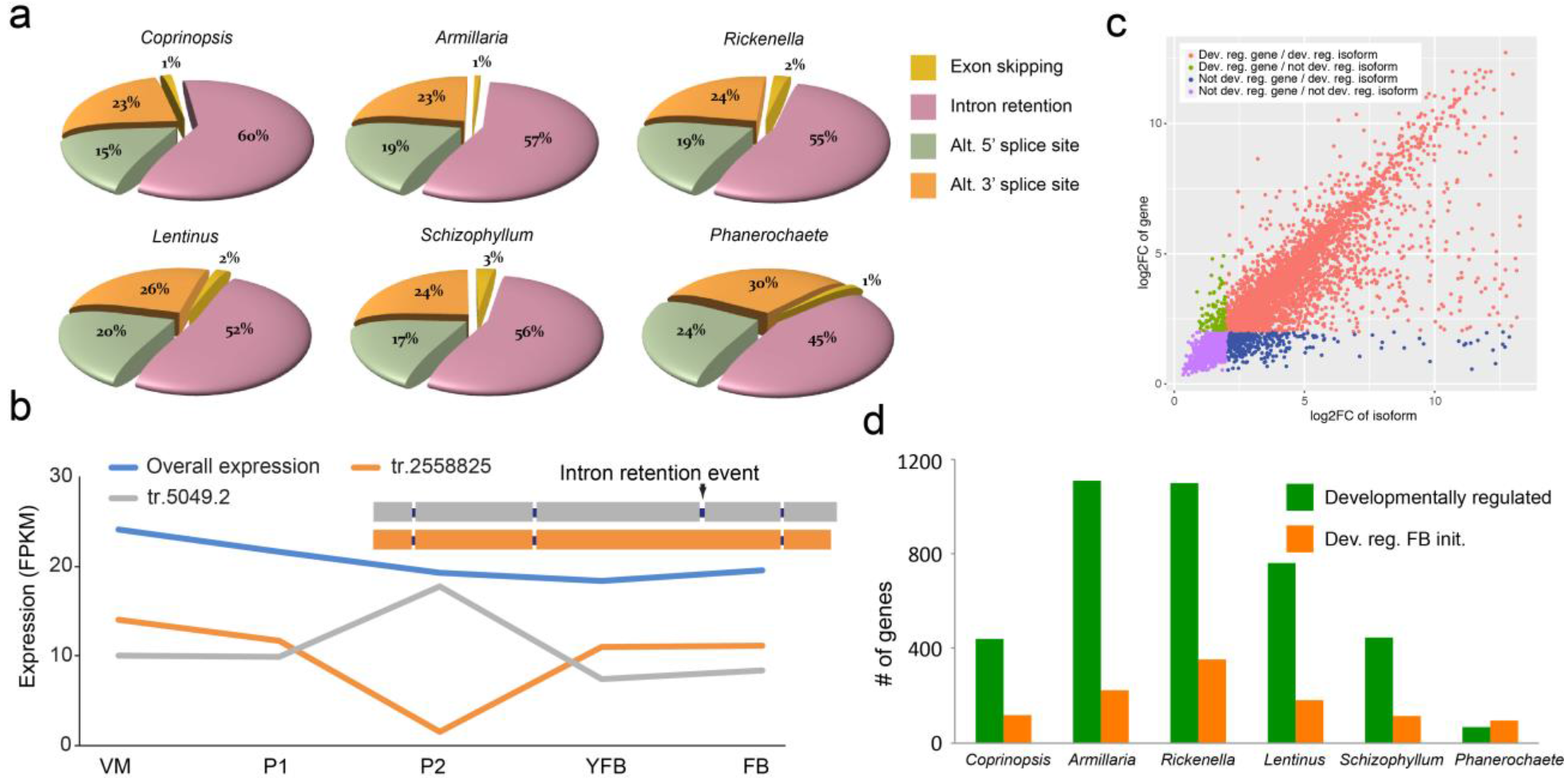
Splicing patterns through fruiting body development. **a**. distribution of four main alternative splicing events across species (see also Supplementary Table 9). **b**. Genes with little dynamics can contain developmentally regulated transcripts, Schco3_2558825 is shown here as an example **c**. Scatterplot of gene versus transcript expression dynamics in *Coprinopsis cinerea*, highlighting developmentally regulated transcripts (red). Plotted are fold change values between the minimum and maximum expression across all developmental stages for genes (y axis) and alternative transcript (x axis). Developmentally regulated transcripts (i.e. those that show FC>4 across any two developmental stages and FPKM>4) of non-developmentally regulated genes (FC<4) are highlighted in red. **d**. bar plot of developmentally regulated transcripts that were detected within not developmentally regulated genes.

### Conserved transcriptomic signatures of mushroom development

Our transcriptome data are particularly suited to detecting shared patterns of gene expression across species. We analyzed common functional signals in the six species by estimating the percent of developmentally regulated genes shared by all or subsets of the species based on Markov clustering^25^ of protein sequences. We found 100 clusters containing developmentally regulated genes from all six species, and 196 in five species (Fig. 1/e, Supplementary Table 10). These are enriched for GO terms related to oxidation-reduction processes, oxidoreductase activity, carbohydrate metabolism, among others, corresponding to a suite of carbohydrate active enzymes. Of the 100 families shared by 6 species, fifteen can be linked to the fungal cell wall (FCW), while the remaining families cover diverse cellular functions such as transmembrane transport (6 families), cytochrome p450s (5 families), targeted protein degradation (5 families), or peptidases (3 families). One hundred and four gene families are shared by five species excluding *Phanerochaete* (Fig 1/e), which comes as no surprise, as this species produces the simple crust-like, fruiting bodies. Besides these highly conserved families, an additional 73 functional groups of genes are developmentally regulated in six or five species, but didn’t group into gene families due to their higher rate of evolution. These include most transcription factors, kinases, aquaporins, certain peptidase families and enzymes of primary carbohydrate metabolism (trehalose and mannitol, Supplementary Figure 8) among others (Supplementary Table 10).

Shared developmentally regulated gene families included a conserved suite of CAZymes active on the main chitin and β-1,3-and β-1,6-glucan polymers as well as minor components of the FCW. These included various glycoside hydrolases (GH), hydrophobins, expansin-like proteins and cerato-platanins, among others. A large suite of β-glucanases, chitinases, laccases, endo-β-1,4-mannanases, α-1,3-mannosidases were developmentally regulated, many of which are also expanded in Agaricomycetes (Table 1, Supplementary Fig 7). The expression of glucan-, chitin-and mannose-active enzymes is consistent with active FCW remodeling during fruiting body formation and recent reports of similar genes upregulated in the fruiting bodies of *Lentinula*^12,13,26^, *Flammulina^27^* and *Coprinopsis^28^.* Kre9/Knh1 homologs are developmentally regulated in all species and are overrepresented in mushroom-forming fungi (P = 1.45×10^−5^, FET). This family is involved in β-glucan assembly in *Saccharomyces* and has putative signaling roles through an interaction with MAP kinases^29^. Although generally linked to cellulose degradation^30,31^, expansins, lytic polysaccharide monooxygenases and cellobiose dehydrogenases have recently been shown to target chitin polymers^32,33^ or are expressed in fruiting bodies of *Pycnoporus*^34^ and *Flammulina*^35^, suggesting a role in fruiting body development. In addition, developmental expression of two alginate lyase-like families (Table 1) were shared by 6 species and that of a β-glucuronidase (GH79_1) was shared by 4 species *(Armillaria, Coprinopsis, Rickenella* and *Lentinus*). While the targets of these families in fruiting bodies are currently unknown, their conserved expression pattern suggests roles in polysaccharide metabolism during development^36^. Comparison across 201 genomes revealed that 24 of these families have undergone expansions in the Agaricomycetes (Table 1, Supplementary Table 3). In summary, CAZymes might be responsible for producing fruiting body-specific FCW architectures, confer adhesive properties to neighboring hyphae or plasticity for growth by cell expansion. We, therefore, suggest that FCW remodeling comprises one of the foundations of the transition to complex multicellularity during the life cycle of fungi.

**Table 1.**
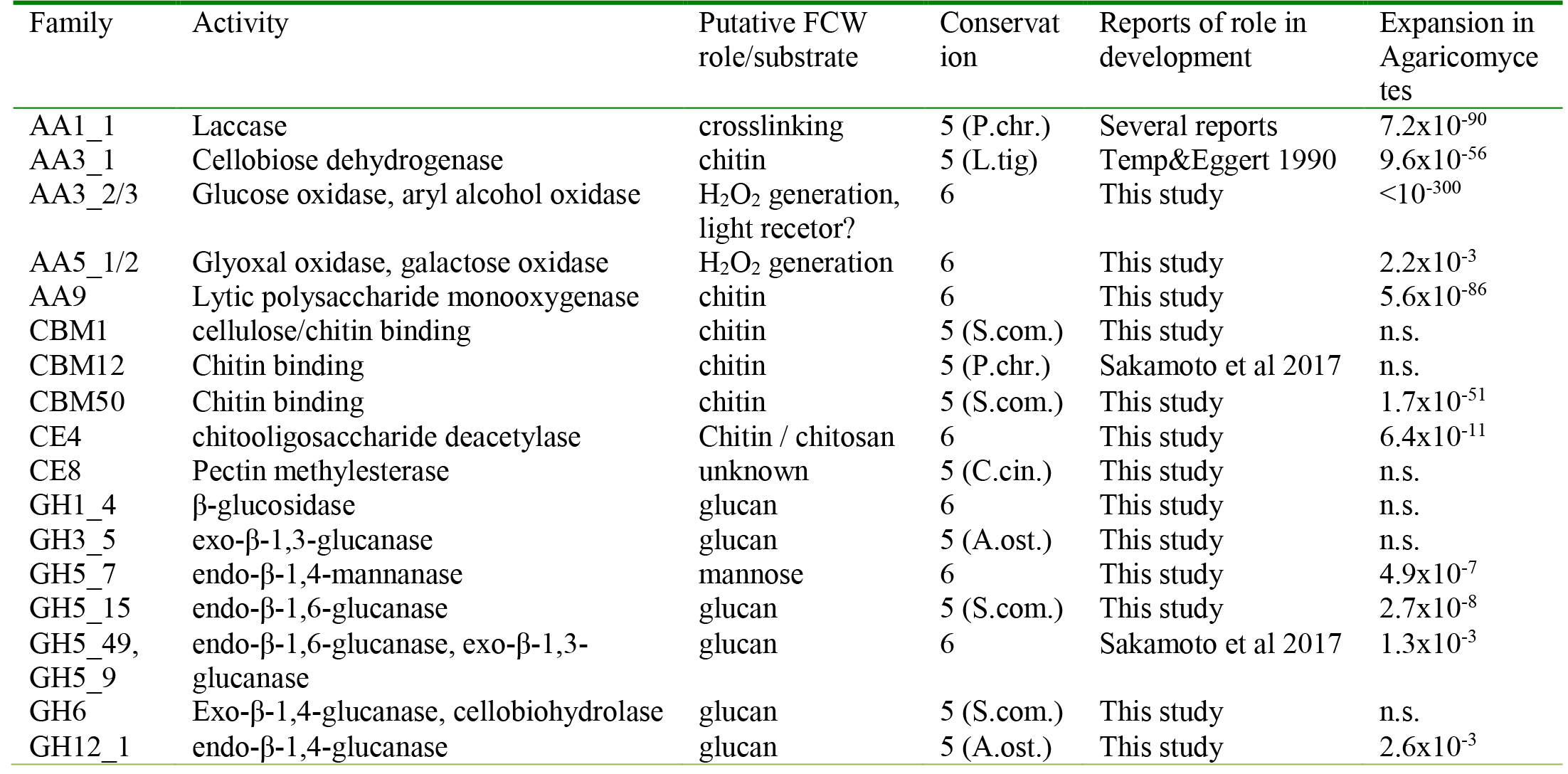
**Conserved developmentally regulated (CAZyme) families and associated modules**

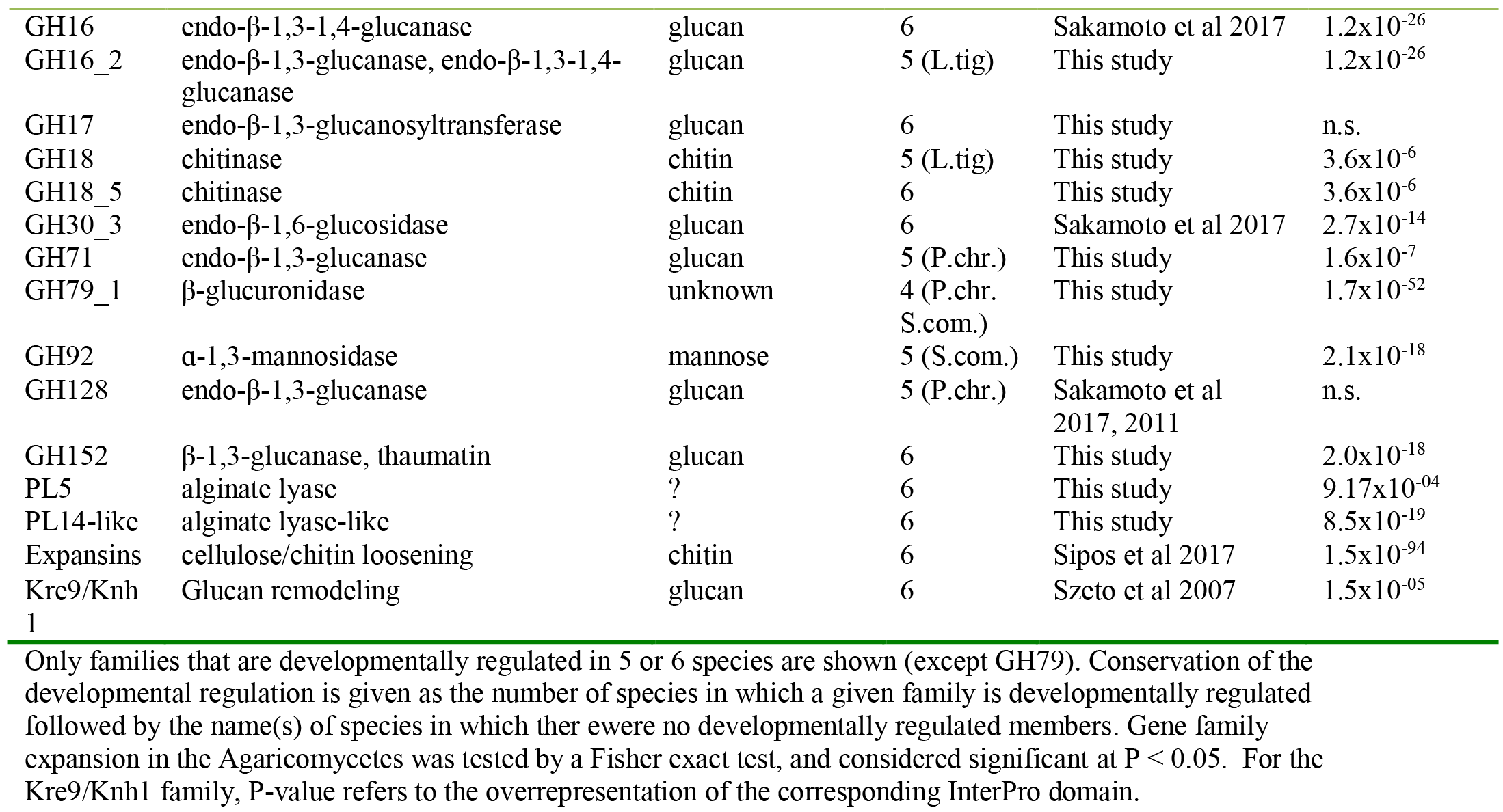

A significant fraction of conserved developmentally regulated genes carry extracellular secretion signals and were predicted to be glycosylphosphatidylinositol (GPI) anchored (Fig 3/a, Supplementary Fig. 9). These include diverse FCW-active proteins, such as laccases (AA1), glucanases (GH5, GH16, Kre9/Knh1 family), chitooligosaccharide deacethylases, but also lectins, A1 aspartic peptidases and sedolisins, among others (Supplementary Table 11). GPI-anchored proteins often mediate adhesion in filamentous and pathogenic fungi^37^, but it is not known whether similar mechanisms are at play in fruiting bodies^7^. Laccases and glucanases could facilitate adhesion by oxidative crosslinking or other covalent modifications of neighboring hyphal surfaces, although more data is needed on the biochemistry involved. Nevertheless, it seems safe to conclude that FCW-active proteins may bind neighboring hyphae through covalent FCW modifications in fruiting bodies, which would represent a unique adhesion mechanism among complex multicellular organisms. Homologs of cadherins (adhesive proteins of animals) are enriched in Agaricomycetes compared to other fungi (*P* = 1.1×10^−4^, FET) and were developmentally regulated in all species. Although fewer in numbers than in animals, their convergent expansion in complex multicellular fungi and metazoans could indicate recurrent co-option for developmental functions.

Fruiting body secretomes contained a rich suite of genes encoding small secreted proteins (SSPs, <300 amino acids, with extracellular secretion signal). Of the 190-477 SSPs predicted in the genomes of the six species, 20-61% are developmentally regulated, with ~20% being conserved across the six species (Fig. 3/b Supplementary Fig. 10). Conserved and annotated genes comprise various FCW-related families, such as hydrophobins, cerato-platanins, cupredoxins, lectins, Kre9/Knh1, GH12 and LysM domain proteins, among others (Fig. 3/c, Supplementary Fig. 11). Hydrophobins and cerato-platanins are SSPs that self-assemble into a rodlet layer on the cell surface, confer hydrophobic surfaces to hyphae that hinder soaking of fruiting bodies with water. They are hypothesized to mediate adhesion, the aeration of fruiting bodies^8,38^, or pathogenicity^39^. As reported previously^8^, most hydrophobin genes are developmentally regulated (Fig. 3/d) and the family is overrepresented in the genomes of mushroom-forming fungi (*P* < 10^−300^, FET). Cerato-platanins are also expanded (*P* = 1.56×10^−50^, FET) and developmentally regulated (except in *Phanerochaete).* In addition to conserved genes, >40% of developmentally regulated SSPs had no functional annotations and/or were species-specific orphans (Fig. 3/c). This proportion is similar to that observed in ectomycorrhiza-induced SSPs^1,40^ and suggests that species-specific secreted proteins have a role also in fruiting body development. Although their function in fruiting bodies is not known, their role in signaling across partners in ectomycorrhizal^40^ and pathogenic interactions^41^, or within species^42,43^, raises the possibility that some of the detected SSPs might act as fruiting body effectors. This could also explain the rich SSP repertoires of saprotrophic Agaricomycetes^44^.

**Fig. 3.**
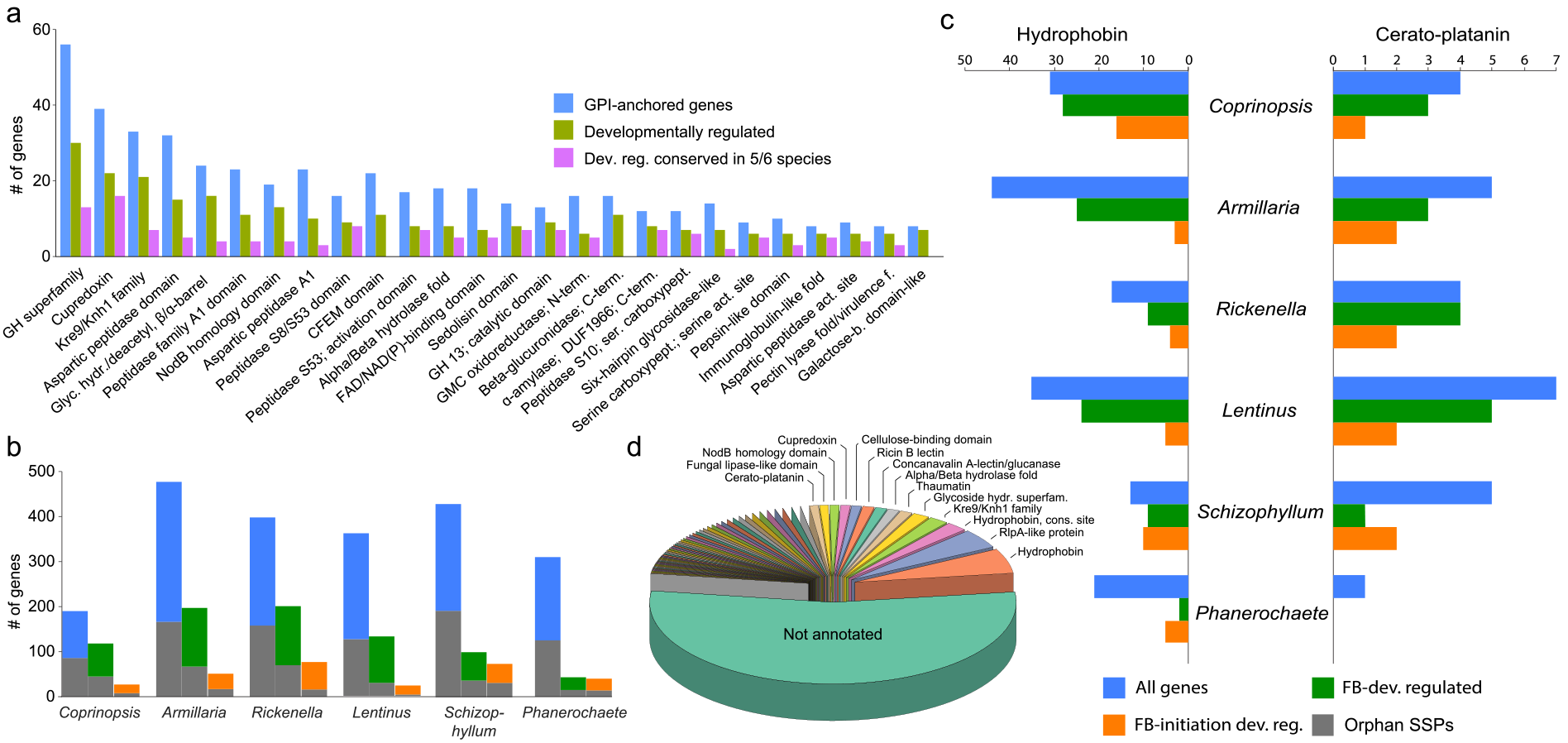
Diverse secreted proteins are developmentally regulated in fruiting bodies. **a**. the distribution of GPI-anchored secreted protein repertoire, its developmentally regulated and conserved developmentally regulated subsets. **b**. Numbers of all and developmentally regulated small secreted proteins, with orphan genes shaded differently. **c**. Copy number distribution and developmental regulation of hydrophobins and cerato-platanins in the six species. **d**. Functional annotation of small secreted proteins in the six species.

### Targeted protein degradation shows striking expansion in mushrooms

We found a strong signal for developmental expression of components the E3 ubiquitin ligase complex. Several genes encoding F-box proteins, RING-type zinc-finger and BTB/POZ domain proteins are developmentally regulated in all species, often displaying tissue or developmental stage-specific expression peaks (Fig. 4, Supplementary Fig. 12). These gene families are also strongly overrepresented in the genomes of mushroom-forming fungi compared to related filamentous fungi and yeasts (Fig 4/a). For example, while yeasts and filamentous fungi possess ~20 and 60-90 F-box proteins^45^, respectively, mushroom-forming fungi have 67-1,199 copies (mean: 274), comparable to the numbers seen in higher plants^46^ and resulting predominantly from recent tandem duplications (Fig. 4/b, Supplementary Fig. 13). They mostly showed a single peak in expression and many of them were upregulated at fruiting body initiation or in caps, gills and stipes (Fig 4/c).The numbers of developmentally regulated F-box, RING-type zinc-finger and BTB/POZ domain containing genes found in the six species show a good correlation with fruiting body complexity but a poor correlation with the number of expressed genes (Fig 4/d), suggesting a link between the expansion of these genes and the evolution of complex fruiting body morphologies. These genes define the target specificity of E3 ubiquitin ligases ^45,47^, which enables a tight regulation of selective proteolysis during development^46^. In plants, F-box proteins can also act as transcriptional regulators^48^, although this is yet to be proven in fungi. On the other hand, ubiquitin conjugating (E2) enzymes are developmentally regulated only in *Coprinopsis, Armillaria* and *Rickenella*, whereas ubiquitin activating (E1) enzymes, cullins, SKP1, HECT-type ubiquitin ligases are neither developmentally regulated nor significantly overrepresented in mushroom-forming fungi (Fig 4/e, P > 0.05, FET). With the exception of *Coprinopsis,* we did not detect specific expression patterns of neddylation and deneddylation genes as reported for the Ascomycota^49^. Taken together, we observed a striking expansion and distinctive expression patterns of genes that define target-specificity of the E3 ubiquitin ligase complex (F-box, RING and BTB/POZ proteins) in Agaricomycetes. This parallels F-box gene expansion in plants which, combined with their widespread role in development^46,50^, suggests that they likely have key roles in complex multicellular development in mushroom-forming fungi.

**Fig. 4.**
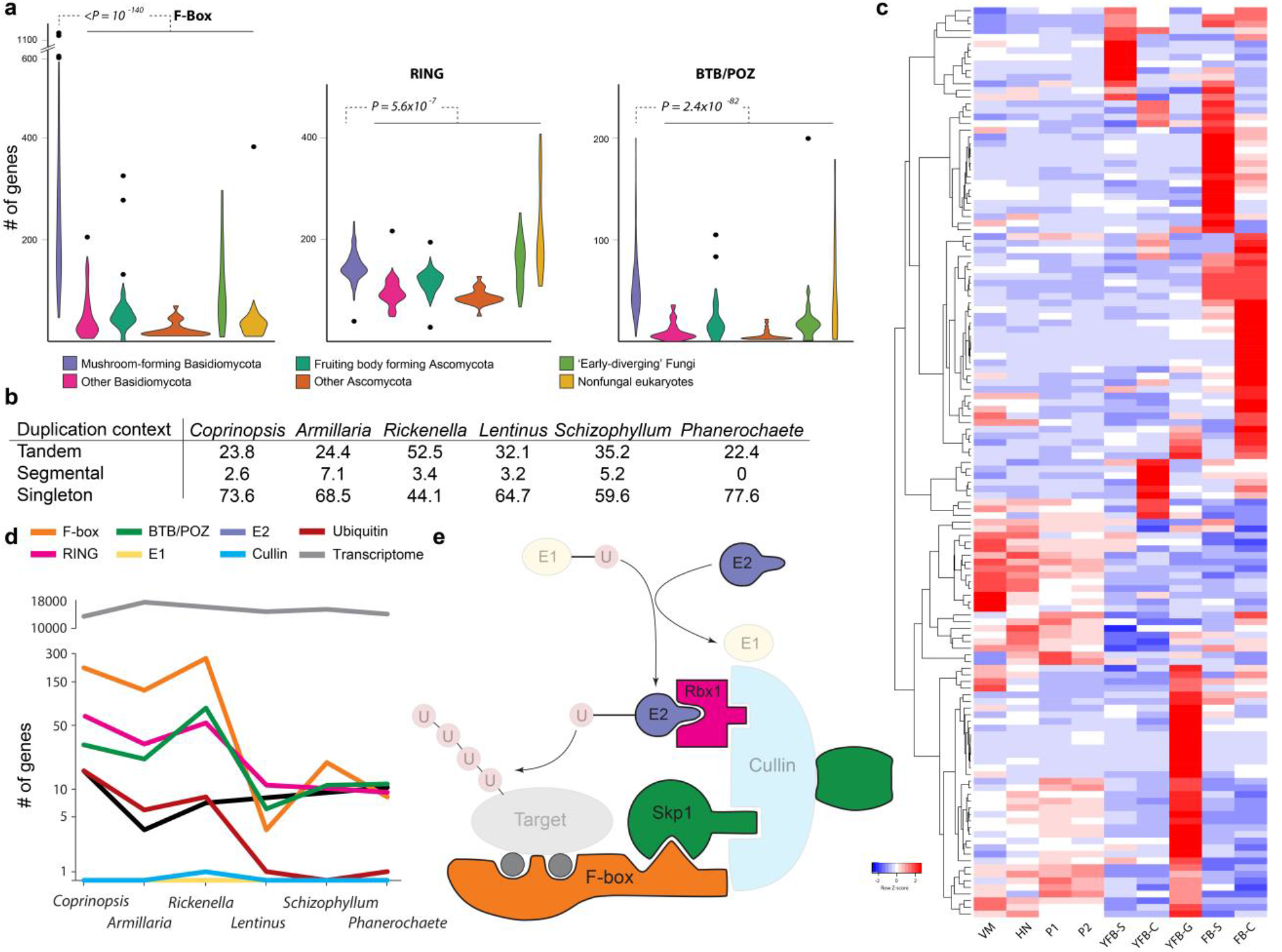
**a**, Copy number distribution of F-box, RING-type zinc finger and BTB/POZ domain proteins showing their enrichment in mushroom-forming fungi compared to other fungal groups. **b**, The context of gene duplications of F-box proteins in the genomes of six analyzed species. **c**, heatmap of developmentally regulated F-box proteins in *Coprinopsis cinerea.* Genes were hierarchically clustered based on gene expression similarity using average linkage clustering. Abbreviations: VM - vegetative mycelium, HN - hyphal knot, P1 - stage 1 primordium, P2 - stage 2 primordium, YFB - young fruiting body, FB - mature fruiting body, ‘-C’ - cap, ‘-S’ - stipe, ‘-G’ - gills. **d**, The correlation between morphological complexity of fruiting bodies and the number of developmentally regulated elements of the E3 ligase complex. **e**, Outline of the E3 ubiquitin ligase complex highlighting members expanded and developmentally regulated (solid) in mushroom-forming fungi. Transparent members are not expanded nor developmentally regulated.

### Key multicellularity-related genes are developmentally regulated in fruiting bodies

Complex multicellularity in fungi is implemented by the reprogramming of hyphal branching patterns, followed by their adhesion and differentiation^7^. This assumes mechanisms for cell-to-cell communication, adhesion, differentiation and defense. We examined the expression dynamics of gene families related to these traits, including transcription factors (TFs), protein kinases, adhesion and defense-related genes. Like other complex multicellular lineages, mushroom-forming fungi make extensive use of transcription factors (TFs) in development. To identify development-related TFs, we manually curated TF candidate genes to exclude ones that nonspecifically bind DNA. The resulting TFomes contain 278-408 genes, of which 4.5-64% were developmentally regulated (Supplementary Fig. 14). These were dominated by C2H2 and Zn(2)C6 fungal type, fungal trans and homeodomain-like TFs (Fig 5/a). Although transcription factor families were usually not conserved, we found 5 TF families that contained developmentally regulated genes from 5 or 6 species (Supplementary Table 10). These included C2H2-type zinc fingers (including *c2h2* of *Schizophyllum*^14,51^), Zn(2)-C6 fungal-type and homeobox TFs (containing *hom1* of *Schizophyllum*^14,51^). Two clusters of C2H2 and homeobox TFs showed expression peaks in stipes of *Coprinopsis, Lentinus, Armillaria* and *Rickenella*, confirming previous reports of *Hom1* expression in *Coprinopsis^52^* and *Schizophyllum^51,53^.* Members of the white collar complex were developmentally regulated in all species except *Phanerochaete*, mostly showing a significant increase in expression at initiation. However, these genes did not group into one family in the clustering, which was a common pattern for TF families, perhaps caused by their high rate of sequence evolution.

**Fig. 5.**
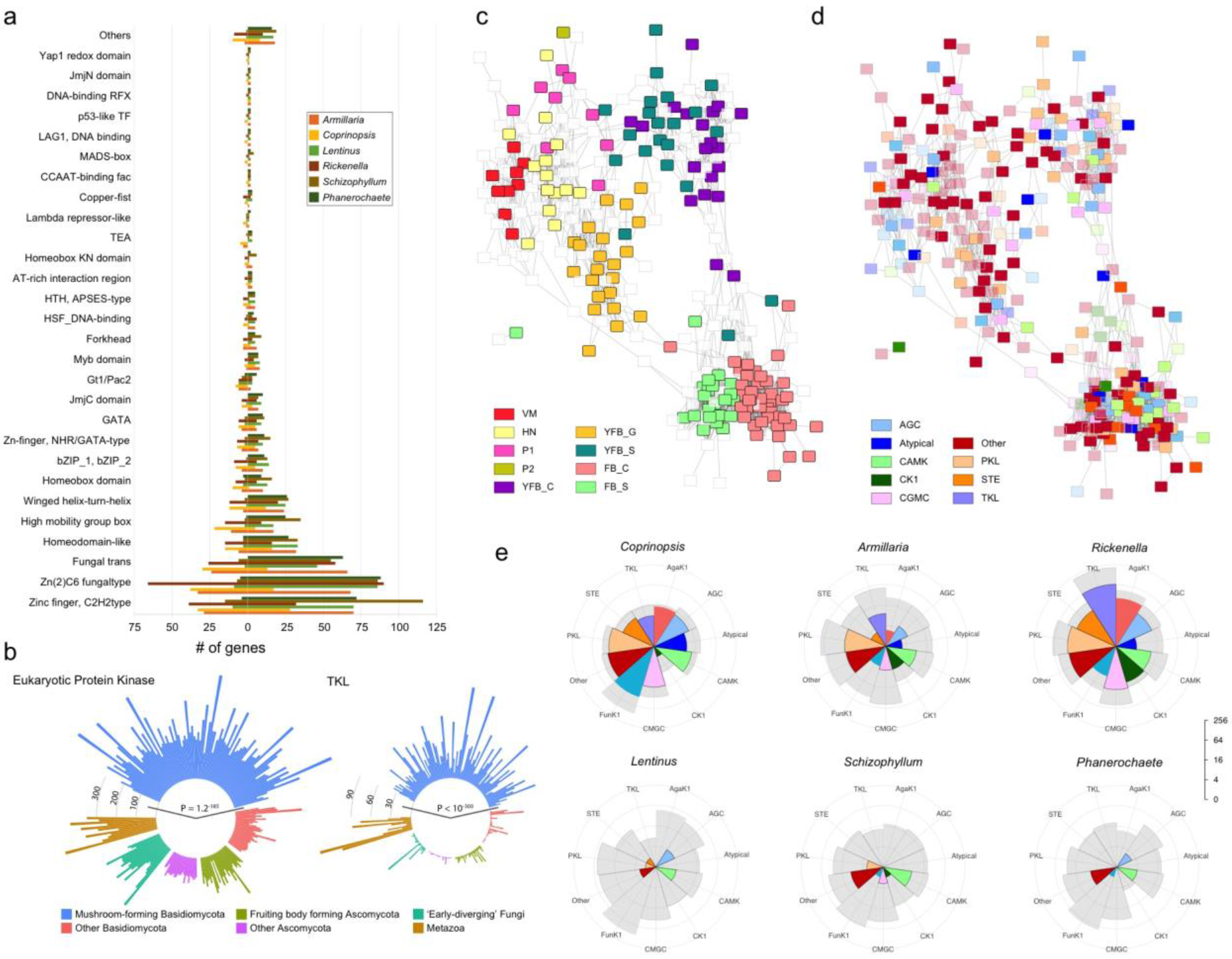
Expression and copy number distribution of transcription factor and kinase genes. **a**. transcription factor family distribution and the proportions of developmentally regulated (left) versus not regulated (right) genes across six species. **b**. Circular bar diagram of eukaryotic protein kinase (left) and tyrosine kinase-like (right) repertoires of mushroom-forming fungi versus in other fungal groups and non-fungal eukaryotes. P values of overrepresentation in mushroom-forming fungi are given for both plots. **c** and **d**. Co-expression network of 306 expressed kinases of *Coprinopsis cinerea* overlaid with expression peak (**c**) and kinase classification (**d**). Pairwise correlation coefficients of expression profile similarity were calculated for each pair of kinase genes and networks were visualized with a correlation coefficient cutoff of 0.825. e. Kinome (grey fill) and developmentally regulated kinome (solid fill) repertoires of the six species split by kinase classification. The families Funk1 and Agak1 of the ‘Other’ group are shown separately. Axes are transformed to log_2_ scale.

Communication among cells by various signaling pathways is paramount to the increase of distinct cell types in evolution. In accordance with the higher complexity of mushroom-forming fungi, their kinomes are significantly larger than that of other fungi (P = 1.1×10^−184^, FET, Fig 5/b), due to expansions of the eukaryotic protein kinase superfamily. We classified protein kinases into nine groups^54^ using published kinome classifications for *Coprinopsis*^3^. The six mushroom kinomes (Supplementary Table 12) have a similar composition, with PKL, CMGC and CAMK families being most diverse and RGC and tyrosine kinases missing. The Agaricomycetes-specific FunK1 family is expanded significantly, as reported earlier^3^. Tyrosine kinase-like kinases also show a strong expansion in Agaricomycetes (P < 10^−300^), consistent with observations in *Laccaria bicolor*^55^ and other species^56^. Histidine kinases are underrepresented (P = 8.95×10^−36^) relative to other fungal groups.

A kinase co-expression network revealed tissue specificity as the main driver of network topology (Fig 5/c-d), with most kinases showing an expression peak late in development. Kinases with early expression peaks are mostly highly expressed through multiple stages, resembling early expressed modules of *Coprinopsis* (Fig. 1/c, Supplementary Fig. 15). Many CAMK family members showed expression peaks in cap and gill tissues. However, overlaying the network with classification shows no general enrichment of any family in developmental stages or tissues (Fig 5/d-e), indicating diverse co-option events for development at the highest level of kinase classification. The FunK1 family has been linked to multicellular development, based on an upregulation in fruiting bodies^3^. In our species FunK1 comprises 4-33% of the kinome and 0.2-35% of the developmentally regulated kinases, although this Fig. resembles that of other kinase families. Members of the highly expanded tyrosine kinase like (TKL) family (Fig 5/e) are developmentally regulated in *Coprinopsis, Armillaria* and *Rickenella*, but not in species with simpler morphologies. The expansion and developmental expression of the TKL family in mushroom-forming fungi is a remarkable case of convergence with complex multicellular plants^57,58^ and animals^58^ and may be related to parallel increases in organismal complexity. Yet, while most plant TKL genes have receptor-like architectures^59^, we found no evidence of extracellular domains or secretion signals in TKL genes of mushroom-forming fungi, suggesting they orchestrate signal transduction via soluble kinases or other mechanisms different from those of multicellular plants and animals.

Fungal immune systems comprise innate chemical defense mechanisms against metazoan predators and bacterial and fungal infections^60^. We catalogued eleven families of defense effector proteins and their expression to assess the conservation of the defensive arsenal of Agaricomycetes. Genomes of mushroom-forming fungi harbor highly species-specific combinations of defense-related genes encoding pore-forming toxins, cerato-platanins, lectins and copsins, among others, with most of them being developmentally expressed and upregulated at fruiting body initiation (Supplementary Fig 16). Of the eleven families only three were conserved, and only one (thaumatins) was developmentally regulated in all six species, which can display either endoglucanase or antimicrobial activity, depending on the structure of the mature protein. *In silico* structure prediction identified an acidic cleft in fruiting body expressed thaumatins (Supplementary Fig 16), consistent with an antimicrobial activity. Several defense-related lectins have been reported from fruiting bodies^61^, although lectins have been implicated in cell adhesion and signaling too. Agaricomycete genomes encode at least 17 lectin and 2 lectin-like families, of which seven are significantly overrepresented (P<0.05, FET, Supplementary Table 3). Developmentally regulated lectins belong to nine families, with 4-7 family per species, but only ricin B lectins were developmentally regulated in all six species (Supplementary Fig 17). Other lectin families show a patchy phylogenetic distribution, which is also reflected in their expression patterns in fruiting bodies. Several lectins are induced at fruiting body initiation, including all previously reported nematotoxic *Coprinopsis* lectins (CCL1-2, CGL1-3 and CGL3). Taken together, the defense effector and lectin-encoding arsenal of mushroom-forming fungi shows a patchy phylogenetic distribution, consistent with high gene turnover rates or gains via horizontal gene transfer^60^. Accordingly, expressed defense gene sets are highly species-specific, with most of the encoded genes upregulated in fruiting bodies, suggesting that chemical defense is a key fruiting body function.

### Discussion

We charted the transcriptomic landscape of multicellular development in 6 phylogenetically diverse mushroom-forming species and performed comparative analyses of >200 genomes. We pinpointed nearly 300 conserved gene families, and another 73 gene groups with developmentally dynamic expression in ≥5 species, as well as 631 domains significantly overrepresented in mushroom-forming fungi. These are enriched in cell wall modifying enzymes, various secreted proteins (including GPI-anchored and small secreted proteins), components of the ubiquitin ligase complex, kinases or transcription factors. Lectins and defense effectors, on the other hand, showed species-specific repertoires, indicating a higher rate of evolutionary turnover. These data provide a framework for elucidating the core genetic program of fruiting body formation and will serve as guideposts for a systems approach to understanding the genetic bases of mushroom development and multicellularity.

Complex multicellularity evolved in five lineages, of which plants, animals and fungi are the most diverse^7,58^. At the broadest level of comparison, these lineages evolved similar solutions to cell adhesion, communication, long-range transport and differentiation^58,62,63^. As in animals and plants, protein kinases, putative adhesive proteins, defense effectors, and certain transcription factors have expanded repertoires in mushroom-forming fungi and show developmentally dynamic expression patterns. Examples for convergent expansions in mushroom-forming fungi, in plants and/or animals include TKL family kinases, F-box proteins and cadherins, indicating that ancient eukaryotic gene families with apt biochemical properties have been repeatedly co-opted for complex multicellularity during evolution. F-box proteins showed the largest expansion in Agaricomycetes across all families and was the largest developmentally regulated family, along with RING-type and BTB domain proteins. Among independently evolved fruiting body forming fungi^7,49,64^, Agaricomycetes share some similarity with the Pezizomycotina (Ascomycota), many of which also produce marcoscopic fruiting bodies. For example, laccases, lectins, several transcription factors and signal transduction systems have also been implicated in fruiting body formation in the Pezizomycotina, although at the moment it is unclear how frequently convergent expansions and co-option events can be observed in independently evolved lineages of complex multicellular fungi^7^.

Mushroom-forming fungi also show several unique solutions for multicellularity, as expected based on their independent evolutionary origin. These are, in part, explained by the very nature of fungi: complex multicellularity comprises the reproductive phase of the life cycle (except in sclerotia, rhizomorphs) and so mechanisms have evolved for sensing when fruiting body formation is optimal (e.g. nutrient availability, light). Mushroom development can be partitioned into an early phase of cell proliferation and differentiation and a growth phase of rapid cell expansion, a division evident on gene co-expression profiles as well. Broadly speaking, this is similar to the development of fleshy plant fruits, although mechanisms are likely to be convergent, like many other aspects of mushroom development.

This work has provided a glimpse into the core genetic toolkit of complex multicellularity in mushroom-forming fungi. Our comparative transcriptomic genomic analyses revealed gene families, most of which reported as novel, with conserved developmental expression in fruiting bodies, with scope to increase the resolution both phylogenetically and among cell types (e.g. by single-cell RNA-Seq). Such data should help defining the conserved genetic programs underlying multicellularity in mushroom-forming fungi, and uncovering the evolutionary origins of a major complex multicellular lineage in the eukaryotes.

## Methods

### Strains, fruiting protocols and nucleic acid extraction

For fruiting *Coprinopsis cinerea* strain #326 (*A43mut B43mut pab 1-1*) an agar disk (5 mm in diameter) was placed on the center of YMG/T agar media (4 g yeast extract, 10 g malt extract, 4 g glucose, 10 g agar media with 100 mg tryptophane added after cooling^65^) at 37 °C for five days in the dark. When the colonies reached the 1-2 mm distance from the edge of Petri dishes they were placed into 25 °C for one week in a 12 hrs light/12 hrs dark cycle for fruiting. Fruiting body stages were defined following standard conventions^5^. Exact alignment of developmental stages across species was impossible, but we made an attempt to define functionally putatively homogeneous stages to follow the general notation of mushroom developmental stages as closely as possible in each species. Nevertheless, the array of sample types differed from species to species, due to the morphological diversity of fruiting bodies or limitations in dissectability. In *Coprinopsis*, vegetative mycelium, hyphal knot, stage 1 and stage 2 primordia, young fruiting body cap, gills and stipe, fruiting body cap and stipe were harvested for RNA extraction. The hyphal knot stage was defined as an up to 0.5 mm diameter condensed hyphal aggregate. Stage 1 primordia were defined as up to 2 mm tall shaft like structures, while stage 2 primordia up to 4 mm tall fruiting body initials with visible differentiation of cap and stipe initials. Young fruiting bodies were up to 15 mm tall with a slightly elongated stipe and immature basidia. Fruiting bodies had fully extended stipes and caps, being in an early autolytic phase.

Before fruiting, *Schizophyllum commune* the H4-8a and H4-8b monokaryons^4^ were grown on MM medium according to Dons et. al.^66^. After dikaryon formation an agar plug (5 mm) was placed on the center of fresh MM medium at 30 °C for five days in the dark, then it was placed at 25 °C for 10 days in a 12/12 hrs light/dark cycle (cool white F18w/840), upside down. Dikaryotic vegetative mycelium, stage 1 and 2 primordia, young fruiting body and fruiting body stages were harvested for RNA-seq. We defined stage 1 primordia as up to 2 mm fruiting body initials, stage 2 primordia as 3-4 mm tall initials with an apical pit on the top, the young fruiting body as a 5-7 mm tall cup-like structure with visible pseudolamellae inside, while fully expanded ones were considered fruiting body.

Vegetative mycelia of *Lentinus tigrinus* RLP-9953-sp were maintained on MEA (20 g malt extract, 0.5 g yeast extract, 15 g agar for 1L). For fruiting a mycelial plug was placed (5 mm diameter) on modified sawdust-rice bran medium^67^ (1 part wheat bran and 2 parts aspen sawdust wetted to 65% moisture for 100 ml in a 250 ml beaker). The culture was incubated for 21 days at 30 °C in the dark, then placed in a moist growth chamber at 25 °C in a 12/12 hour light/dark cycle. Vegetative mycelia, stage 1 primordia, stage 2 primordia cap and stipe, young fruiting body cap and stipe and fruiting body cap and stipe tissues were harvested for RNA-Seq. Stage 1 primordium was defined as a 5-20 mm tall white stalk-structure without any differentiation of a cap initial, stage 2 primordium was defined as a 15 - 25 mm tall stalk-like structure with a brown apical pigmentation (cap initial), young fruiting body had and up to 5 mm wide brown cap initial with just barely visible gills on the bottom, growing on a 30-40 mm tall stipe, fruiting body was 50-70 mm tall, with fully flattened (but not funnel-shaped) cap.

*Phanerochaete chrysosporium* RP-78 was fruited on YMPG media (10 g glucose, 10 g malt extract, 2 g peptone, 2 g yeast extract, 1 g asparagine, 2 g KH_2_PO4, 1 g MgSO_4_ x 7 H_2_O, 20 g agar for 1L with 1 mg thiamine added after cooling) covered with cellophane for 7 days at 37 °C in the dark, then placed in a moist growth chamber at 25 °C in an area with dimmed ambient light conditions, following the recommendations of Jill Gaskell (US Forest Products Laboratory, Washington, D. C., USA). Vegetative mycelium, young fruiting body and fruiting body stages were harvested for RNA extraction. Young fruiting body stage was defined as fruiting body initials that forms a compact mat well-delimited from the surrounding vegetative mycelium, while the fruiting bodies were harvested just after they started releasing spores (visible on the lids of Petri dishes).

*Rickenella mellea* SZMC22713 was cultured on Fries Agar^68^ for harvesting vegetative mycelium for RNA and DNA extraction. DNA for genome sequencing was extracted using the Blood & Cell Culture DNA Maxi Kit (Qiagen) from 300 mg finely ground mycelium powder according to the manufacturer’s instructions. The internal transcribed spacer region was PCR amplified and sequenced to verify strain identity. For RNA-Seq, fruiting body stages were collected in November 2016 from Kistelek, Hungary (approx. coordinates: 46.546309, 19.954507). Stage 1 primordium was defined as an approximately 1 mm tall, shaft-like, pear-shaped structure, without any visible cap initial, stage 2 primordium was described as a 2-3 mm tall structure with a small cap initial, young fruiting body was defined by the 5-15 mm tall structure with a 1-2 mm wide cap, and the fruiting body was characterized by a fully expanded cap on the top of a 20-32 mm tall stipe.

Data for *Armillaria ostoyae* C18/19 were taken from our previous study^16^, with the following stages defined: vegetative mycelium, stage 1 primordium, stage 2 primordium cap and stipe, young fruiting body cap and stipe, and fruiting body cap, stipe and gills.

For RNA extraction all samples were immediately placed on liquid nitrogen after harvesting and stored at −80 °C until use. Frozen tissues were weighed and 10-20 mg of *C. cinerea, S. commune, P. chrysosporium* and *R. mellea* and 50-75 mg of *L. tigrinus* were transferred to a pre-chilled mortar and ground to a fine powder using liquid nitrogen. We extracted RNA of *C. cinerea, S. commune, P. chrysosporium* and *R. mellea* using the Quick-RNA™ Miniprep (Zymo Research), or the RNeasy Midi Kit (QIAGEN) for *L. tigrinus.* Both of the kits were used according to the manufacturer’s instructions.

### De novo draft genome for *Rickenella mellea*

The genome and transcriptome of *Rickenella mellea* were sequenced using Illumina platform. The genomes were sequenced as pairs of Illumina standard and Nextera long mate-pair (LMP) libraries. For the Illumina Regular Fragment library, 100 ng of DNA was sheared to 300 bp using the Covaris LE220 and size selected using SPRI beads (Beckman Coulter). The fragments were treated with end-repair, A-tailing, and ligation of Illumina compatible adapters (IDT, Inc) using the KAPA-Illumina library creation kit (KAPA biosystems).

For the Illumina Regular Long-mate Pair library (LMP), 5 ug of DNA was sheared using the Covaris g-TUBE(TM) and gel size selected for 4 kb. The sheared DNA was treated with end repair and ligated with biotinylated adapters containing *loxP*. The adapter ligated DNA fragments were circularized via recombination by a Cre excision reaction (NEB). The circularized DNA templates were then randomly sheared using the Covaris LE220 (Covaris). The sheared fragments were treated with end repair and A-tailing using the KAPA-Illumina library creation kit (KAPA biosystems) followed by immobilization of mate pair fragments on streptavidin beads (Invitrogen). Illumina compatible adapters (IDT, Inc) were ligated to the mate pair fragments and 8 cycles of PCR was used to enrich for the final library (KAPA Biosystems).

Stranded cDNA libraries were generated using the Illumina Truseq Stranded RNA LT kit. mRNA was purified from 1 μg of total RNA using magnetic beads containing poly-T oligos.
mRNA was fragmented and reversed transcribed using random hexamers and SSII (Invitrogen) followed by second strand synthesis. The fragmented cDNA was treated with end-pair, A-tailing, adapter ligation, and 8 cycles of PCR.

The prepared libraries were quantified using KAPA Biosystem’s next-generation sequencing library qPCR kit and run on a Roche LightCycler 480 real-time PCR instrument. The quantified libraries were then multiplexed with other libraries, and the pool of libraries was then prepared for sequencing on the Illumina HiSeq sequencing platform utilizing a TruSeq paired-end cluster kit, v4, and Illumina’s cBot instrument to generate a clustered flow cell for sequencing. Sequencing of the flow cell was performed on the Illumina HiSeq2500 sequencer using HiSeq TruSeq SBS sequencing kits, v4, following a 2×150 indexed run recipe (2×100bp for LMP).

Genomic reads from both libraries were QC filtered for artifact/process contamination and assembled together with AllPathsLG v. R49403^69^. Illumina reads of stranded RNA-seq data were used as input for de novo assembly of RNA contigs, assembled into consensus sequences using Rnnotator (v. 3.4)^70^. Both genomes were annotated using the JGI Annotation Pipeline and made available via the JGI fungal portal MycoCosm^71^. Genome assemblies and annotation were also deposited at DDBJ/EMBL/GenBank under the accession XXXXX (TO BE PROVIDED UPON PUBLICATION).

### RNA-Seq

Whole transcriptome sequencing was performed using the TrueSeq RNA Library Preparation Kit v2 (Illumina) according to the manufacturer’s instructions. Briefly, RNA quality and quantity measurements were performed using RNA ScreenTape and Reagents on TapeStation (all from Agilent) and Qubit (ThermoFisher); only high quality (RIN >8.0) total RNA samples were processed. Next, RNA was DNaseI (ThermoFisher) treated and the mRNA was purified and fragmented. First strand cDNA synthesis was performed using SuperScript II (ThermoFisher) followed by second strand cDNA synthesis, end repair, 3’-end adenylation, adapter ligation and PCR amplification. All of the purification steps were performed using AmPureXP Beads (Backman Coulter). Final libraries were quality checked using D1000 ScreenTape and Reagents on TapeStation (all from Agilent). Concentration of each library was determined using either the QPCR Quantification Kit for Illumina (Agilent) or the KAPA Library Quantification Kit for Illumina (KAPA Biosystems). Sequencing was performed on Illumina instruments using the HiSeq SBS Kit v4 250 cycles kit (Illumina) or the NextSeq 500/550 High Output Kit v2 300 cycles (Illumina) generating >20 million clusters for each sample.

### Bioinformatic analyses of RNA-Seq data

RNA-Seq analyses were carried out as reported earlier^16^. Paired-end Illumina (HiSeq, NextSeq) reads were quality trimmed using the CLC Genomics Workbench tool version 9.5.2 (CLC Bio/Qiagen) removing ambiguous nucleotides as well as any low quality read end parts. Quality cutoff value (error probability) was set to 0.05, corresponding to a Phred score of 13. Trimmed reads containing at least 40 bases were mapped using the RNA-Seq Analysis 2.1 package in CLC requiring at least 80% sequence identity over at least 80% of the read lengths; strand specificity was omitted. List of reference sequences is provided as Supplementary table 1. Reads with less than 30 equally scoring mapping positions were mapped to all possible locations while reads with more than 30 potential mapping positions were considered as uninformative repeat reads and were removed from the analysis.

“Total gene read” RNA-Seq count data was imported from CLC into R version 3.0.2. Genes were filtered based on their expression levels keeping only those features that were detected by at least five mapped reads in at least 25% of the samples included in the study. Subsequently, “calcNormFactors” from “edgeR” version 3.4.2^72^ was used to perform data scaling based on the “trimmed mean of M-values” (TMM) method^73^. Log transformation was carried out by the “voom” function of the “limma” package version 3.18.13^74^. Linear modeling, empirical Bayes moderation as well as the calculation of differentially expressed genes were carried out using “limma”. Genes showing at least four-fold gene expression change with an FDR value below 0.05 were considered as significant. Multi-dimensional scaling (“plotMDS” function in edgeR) was applied to visually summarize gene expression profiles revealing similarities between samples. In addition, unsupervised cluster analysis with Euclidean distance calculation and complete-linkage clustering was carried out on the normalized data using “heatmap.2” function from R package “gplots”.

### Identification of developmentally regulated genes

We considered genes with a Fragments Per Kilobase Million (FPKM) value >1 to have a non-zero expression. Because differentially expressed genes can only be defined in pairwise comparisons of samples and thus didn’t suit our developmental series data, we opted to use the concept of developmentally regulated gene. These were defined as any gene showing an over four-fold change in expression between any two developmental stages or tissue types and a maximum expression level of FPKM > 4 in at least one developmental stage. Comparisons between tissue types were only performed within the respective developmental stage. We distinguished developmentally regulated genes that showed over four-fold upregulation at fruiting body initiation (‘FB-init’ genes) and those that show over four-fold expression dynamics (up-or downregulation) across the range of fruiting body stages (‘FB genes’, i.e. vegetative mycelium excluded). Note that this strategy excludes genes showing highest expression in vegetative mycelium and no dynamics later on, to remove genes with a significant downregulation at the onset of fruiting body development (e.g. those involved in nutrient acquisition by the mycelium).

### Comparative genomic approaches

To obtain characteristic Interpro domain signatures of Agaricomycetes, we assembled a dataset comprising genomes of 201 species; ranging from simple unicellular yeasts to filamentous and complex multicellular fungi. InterProscan version 5.24-63.0 was used to perform IPR searches. The 201 species were categorized into two major groups; mushroom-forming fungi (113 species) and all other fungi (88 species, 1 Cryptomycota, 2 Microsporidia, 2 Neocallimastigomycota, 3 Chytridiomycota, 2 Blastocladiomycota, 14 Zygomycota, 1 Glomeromycota, 38 Ascomycota, 20 non-fruiting body forming Basidiomycota). The enrichment of IPR domains was tested using Fisher’s exact test and corrected for multiple testing by the Benjamini-Hochberg method in R (R core team 2016). P<0.01 was considered significant. Significantly overrepresented IPR domains were characterized by Gene Ontology Terms using IPR2GO.

An all-versus-all protein BLAST was performed for the six species (*A. ostoyae, C. cinerea, S. commune, L. tigrinus, P. chrysosporium, R. mellea*) and for the 201-species dataset using mpiBLAST (v.1.6.0) with default parameters. Clustering was done using Markov Cluster with an inflation parameter of 2.0^25^.

### Reconstruction of alternative splicing patterns

We reconstructed patterns of alternative splicing using the RNA-Seq data for all six species. To this end, we used region-restricted probabilistic modelling (RRPM)^21^ to discover alternative transcripts, as described by Gehrmann et al., Briefly, the genome was split at gene boundaries into fragments, then all RNA-Seq reads were aligned to these fragments with STAR v2.5.3a^75^, in two rounds. The first round of read alignment was run to produce a novel splice junction database, which was used to improve mapping in the second round. Using the BAM file from this alignment, Cufflinks v2.2.1^76^ was run in RABT mode to predict novel transcripts. To restore the context, these sets of transcripts were projected back onto the original annotation. The resulting annotation file was filtered to remove predicted transcripts with no detectable expression (FPKM = 0) or did not have reads supporting its splice junctions. We performed read alignment using STAR again with the same two round method and the new, corrected annotation file and used the Cufflinks suite to estimate the expression level for each transcript. We then aligned reads of each RNA-Seq replicate separately to the genome with updated gene annotation. This resulted in an expression profile for each alternatively spliced transcript, in every developmental stage. We subsequently identified developmentally regulated transcripts using the same functions as described above for genes. For splicing event discovery, we used the ASpli^77^ R package where we used the most significant transcript (the most abundant transcript through the developmental stages) as the reference for event discovery. Custom scripts were used to extract stage and tissue-type specificity and distribution of spliced genes and splicing events.

### Phylostratigraphic analysis

To examine the evolutionary origin of developmentally regulated genes in each species, a phylostatigraphic analysis was performed^19^. First, we assembled a database containing genomes covering the evolutionary route from the most recent common ancestor of cellular organisms to the respective species, by complementing the database of Drost et al.^20^. Fungal, microsporidia and plant genomes were removed from this database and substituted by 416 fungal genomes (all published), including 382 belonging to the Dikarya and 116 to the Agaricomycetes. In addition, 6 microsporidia, 59 plant and 6 Opisthokonta^78^ genomes were added, resulting in a database of 4,483 genomes. The database was divided into age categories (‘phylostrata’) based on the tree available at Mycocosm^71^ and the eukaryotic tree published by Torruella et al.^78^. The oldest phylostratum consisted of bacteria and archaea. Whole proteomes of *Coprinopsis cinerea, Armillaria ostoyae, Schizophyllum commune, Lentinus tigrinus, Phanerochaete chrysosporium* and *Rickenella mellea* were blasted against this database using mpiblast 1.6.0^79^ with default settings. Blast hits were filtered with an E-value cut-off of 1×10^−6^ and a query coverage cut-off of 80%. After filtering, the age of each gene was defined as the node of the tree representing the last common ancestor of the species sharing homologs of the gene, at the specified blast cutoff.

To infer what Agaricomycete-specific genes are preferentially developmentally regulated, we analyzed the enrichment of annotation terms among developmentally regulated genes specific to Agaricomycetes compared to developmentally regulated genes whose origin predates the Agaricomycetes. To this end, we divided the phylostratigraphy profiles into two groups, corresponding to genes that originated before and those that originated after the origin of mushroom-forming fungi (Phylostratum 18). We tested for significant enrichment of IPR domains (evalue < 1e-5) in developmentally regulated genes that originated within the Agaricomycetes, relative to the other group of more ancient developmentally regulated genes using Fisher’s exact test (P < 0.05).

### CAZyme annotation

Genes encoding putative carbohydrate-active enzymes were annotated using the CAZy pipeline. BLAST and Hmmer searches were conducted against sequence libraries and HMM profiles in the CAZy database^80^ (http://www.cazy.org). Positive hits were validated manually and assigned a family and subfamily classification across Glycoside Hydrolase (GH), Carbohydrate Esterase (CE), Glycoside Transferase (GT), Polysaccharide Lyase (PL), Carbohydrate-Binding Module (CBM) and Auxiliary redox enzyme (AA) classes of the CAZy system^81^. Activities were determined by BLAST searches against biochemically characterized subsets of the CAZy database.

### Coexpression analysis

Developmentally regulated genes in each species were clustered into co-expression modules based on their expression dynamics by using the clustering method implemented in Short Time-series Expression Miner (STEM v. 1.3.11)^18,82^. Default parameters were used, except minimum absolute expression change, which was set to 4. Functional annotations of modules were obtained by GO enrichment analyses in TopGO (see below). For a higher-level grouping of coexpression modules, we defined six categories corresponding to early and late expressed genes, cap, stipe and gill specific genes and a mixed category. Coexpression modules were placed in one of these categories if more than half of the module’s members had the same tissue-or stage-specific expression peaks. Modules without stage or tissue specificity were grouped in the mixed category. The early expressed category included coexpression modules with expression peaks in H, P1 or P2 stages, while late module category consisted of modules with young fruiting body and fruiting body stage specific expression peaks. We functionally annotated the modules and higher categories using InterPro Scan v5.24-63.0.

To visualize the kinase expression network across various kinase groups and developmental stages, a co-expression network was visualized using Cytoscape *v*3.6.1 based on pairwise Pearson correlation coefficients for kinase expression patterns in *Coprinopsis cinerea.* Pairwise Pearson correlations coefficients for each kinase gene pair were calculated and a 0.85 cut-off was applied for network construction.

### Functional annotations, GO and Interpro enrichment

Gene Ontology (GO) enrichment analyses were carried out for developmentally regulated genes. For this, we annotated genes with GO terms based on their InterPro domain contents. Analyses were performed using Fisher’s exact test with threshold P<0.05 in the R package topGO. The parameter algorithm weighted01 was chosen. Heatmaps were created using the heatmap.2 function of the R package ‘gplots’. Unsupervised cluster analysis with Pearson’s distance calculation and averaged-linkage clustering was carried out on the FPKM values, and heatmaps was visualised using z-score normalization on the rows via the heatmap.2 function.

Prediction of glycosylphosphatidylinositol anchored proteins (GPI-Ap) for the six species was performed using the portable version of Pred-GPI^83^. From the proteins with a predicted GPI-anchor, we excluded ones which had no extracellular signal sequence, as assessed by SignalP version 4.1^84^. Prediction of Small Secreted Proteins (SSP) for the six species was performed using a modified version of the bioinformatic pipeline of Pellegrin et al.^44^. Proteins shorter than 300 amino acids were subjected to signal peptide prediction in SignalP (version 4.1) with the option “eukaryotic”. Extracellular localisation of these proteins was checked with WoLFPsort version 0.2^85^ using the option “fungi”. Proteins containing transmembrane helix not overlapping with the signal peptide were also excluded. Prediction of transmembrane helices was performed with TMHMM (version 2.0)^86^. Finally, proteins containing a KDEL motif (Lys-Asp-Glu-Leu) in the C-terminal region (prosite accession “PS00014”) responsible for retention in the endoplasmic reticulum (ER) lumen, were identified using PS-SCAN (http://www.hpa-bioinfotools.org.uk/cgi-bin/psscan/psscanCGI.pl)fig and excluded.

We identified transcription factors based on the presence of InterPro domains with sequence-specific DNA-binding activity retrieved from literature data^87,88^ and manual curation of the Interpro-database. Annotated genes were then filtered based on their domain architecture in order to discard genes encoding DNA-binding proteins with functions other than transcription regulation (such as DNA-repair, DNA-replication, translation, meiosis).

We extracted the putative kinase genes from the 6 species based on their InterPro domain composition, and manually curated the classical kinases by excluding domains which correspond to metabolism related kinases and other non-classical protein kinases. The set of proteins having kinase related domains (Supplementary table 12) were subjected to BLAST searches (*BLAST 2.7.1*+, E-value 0.001) against the kinome of *Coprinopsis cinerea*^3^ downloaded from Kinbase (www.kinase.com). The best hits for the six species were classified into eukaryotic protein kinase (ePK) and atypical protein kinases (aPK) and their families and subfamilies as described in the hierarchical kinase classification system^54^.

## Acknowledgements

The authors thank Daniel Cullen and Jill Gaskell (USDA, USA) for the strain of *Phanerochaete* used in this study. This work was supported of the Momentum Program of the Hungarian Academy of Sciences (Contract No: LP2014/12), and of the National Research, Development and Innovation Office (Grant No: GINOP-2.3.2-15-2016-00001). This project has received funding from the European Research Council (ERC) under the European Union’s Horizon 2020 research and innovation programme (grant agreement number 758161 and 716132). The work by the U.S. Department of Energy Joint Genome Institute, a DOE Office of Science User Facility, is supported by the Office of Science of the U.S. Department of Energy under Contract No. DE-AC02-05CH11231.

## Author contributions

K.K. and L.G.N. conceived the study. K.K., B.K., E.A. performed fruiting experiments, RNA isolation and data analysis. B.B., I.N., J.C. obtained and analysed RNA-Seq data. Z.M., N.S., M.V., T.K. and L.G.N. performed comparative genomic and phylogenomic analyses. S.M. analyzed the genomic context of F-box gene duplications. B.H. analyzed CAZymes. E.A. and R.A.O. analyzed transcription factors. B.H. contributed valuable analytical insights. K.B., J.J., A.L., J.P., J.Y. Y.X. and I.V.G. sequences, assembled and annotated the genome of *Rickenella mellea.* L.G.N, K.K., B.B., and U.K. wrote the manuscript. D.S.H. contributed the genome of *Lentinus tigrinus* and analytical insights. All authors read and commented on the manuscript.

## Data availability

Genome assembly and annotation of *Rickenella mellea* was deposited at DDBJ/ EMBL/GenBank under the accession XXXXXX (to be provided upon publication). A Gene Expression Omnibus (GEO) archive of the sequenced *A. ostoyae* libraries was deposited in the NCBI’s GEO Archive at http://www.ncbi.nlm.nih.gov/geo under accession SRP109671.

